# Probing the limited oculomotor range of common marmosets (*Callithrix jacchus*)

**DOI:** 10.1101/2025.11.26.690791

**Authors:** Oviya Mohan, Alexia Nelms, Amy Bucklaew, Dora Biro, Jude F. Mitchell

## Abstract

The ability to scan visual scenes to gather information is a critical adaptive skill across primates. The common marmoset, a small-bodied New World monkey and emerging model of social and visual neuroscience, relies heavily on rapid head movements in addition to eye movements to orient (Pandey et al., 2020; Singh et al., 2025). Previous studies have found a relatively restricted oculomotor range extending out about 10 visual degrees from the central position of rest (Mitchell et al., 2014; Singh et al., 2025). However, previous studies have either used empty arenas for exploration (Singh et al., 2025), or in head-fixed animals used exploration of images potentially biased centrally with posed stimuli (Mitchell et al., 2014). This leaves open questions about whether this restricted oculomotor range is due to physical constraints or a lack of attention-drawing stimuli in the periphery. Understanding marmosets’ oculomotor range is important for applying modern methods that use marker-less pose tracking of the head as a proxy for gaze direction, under the assumption eye gaze is relatively restricted and can be ignored (Meisner et al., 2025). Using high-precision eye tracking and free-viewing of a natural image and video with objects of interest placed in the periphery, we quantified the oculomotor range of head-fixed marmosets. Our results show limited changes in the range reported from previous studies, even with naturalistic stimuli including moving animals that were optimized to encourage peripheral viewing.

## Introduction

It is well established that humans and non-human primates scan visual scenes to gather information about their environment, which is then used to make critical decisions for survival (e.g., Kano & Tomonaga, 2009; Mitchell et al., 2014). The mechanisms by which different primate species achieve this depend on their physiology, anatomy, and the environment in which they evolved, and hence show considerable variation. For example, humans use head and eye movements to orient to objects of interest, with eye movements ranging out to 55 degrees from the central position of rest (Guitton & Volle, 1987) and in macaques ranging out to 30 degrees (Tehovnik et al., 2021). These objects of interest can include food, predators, or even conspecifics and interactions between conspecifics to obtain social information.

In contrast, some smaller New World monkeys, like the common marmoset (*Callithrix jacchus)* and the squirrel monkey (*Saimiri sciureus*), typically use rapid head movements to orient their attention (J. Burkart & Heschl, 2006; J. M. Burkart & Heschl, 2007; McCrea & Gdowski, 2003; Pandey et al., 2020). This is largely because their lighter heads have less inertia (Heiney & Blazquez, 2011), which may enable them to rely more on these head movements for rapid shifts of gaze. However, the extent to which they can move their eyes from the position of rest, i.e., their oculomotor range, has only been quantified under relatively limited experimental conditions. Previous work (Mitchell et al., 2014) demonstrated a restricted oculomotor range of approximately 10 dva (degrees of visual angle) in head-fixed marmosets viewing natural images, compared to macaques in a similar setup, who exhibited a range of up to 20-25 dva. The study used RGB images of natural scenes obtained from the internet that may have been biased towards centrally located objects of interest and thus may not have prompted the marmosets to make eye movements beyond this limited range. Likewise, a recent study using head-mounted eye tracking in freely moving marmosets also reported a similar restricted range (Singh et al., 2025) but only tested single animals in open arenas with no social stimuli or other moving animals. Thus, an open question remains as to whether the previously observed restricted range reflects a strong physical constraint or instead could reflect a lack of objects of interest in the periphery to draw gaze outside of this range.

Marmosets have become an increasingly popular model in studies of visual neuroscience (Mitchell et al., 2014) as well as the neuroscience of social behavior (e.g., J. M. Burkart & Finkenwirth, 2015; Miller et al., 2016; Meisner et al., 2025). This growing interest has made it important to study their behavior in naturalistic contexts where they are able to interact freely with conspecifics. Tracking their visual attention during these interactions remains a crucial method for studying their underlying socio-cognitive abilities (e.g., Meisner et al., 2025). Head-free gaze tracking utilizing marker-less pose estimation and computer vision tools, where the position of the head is used to estimate the gaze of the marmosets, has become a key tool to enable such naturalistic studies (e.g., Xing et al., 2024). Crucially, these head-gaze estimates rely on assumptions about the limited oculomotor range of these monkeys, making it imperative to quantify and verify this range.

In this paper, we used naturalistic stimuli with the aim to create conditions in which animals would be encouraged to look outside the center and motivate the monkeys to their physical oculomotor limit. We recorded videos from the viewpoint of marmoset monkeys inside the colony where they were raised, including neighbors and surrounding families interacting over the extent of the video, thus creating stimuli natural to this set of animals’ upbringing and daily life. We used high-precision eye tracking of head-fixed marmosets viewing these videos to test whether the oculomotor range identified in previous work is as restricted as it appears or instead may depend on the type of stimuli. We also include still images sampled from the video in order to compare to the previous free-viewing study (Mitchell et al., 2014). This will inform us how much of this restricted range is a physical constraint and how much depends on the stimulus.

## Methods

### Subjects

All experimental protocols were approved by the University of Rochester Institutional Animal Care and Use Committee and were conducted in compliance with the National Institute of Health’s guidelines for animal research. Two adult common marmosets (*Callithrix jacchus*), Marmoset A (male) and Marmoset B (female), were used for eye tracking experiments to measure oculomotor range. Subjects were pair-housed at the University of Rochester with a circadian cycle of 12-hour light and 12-hour dark. Both subjects had full access to food and water. They were surgically implanted with head caps to stabilize them for head-fixed eye tracking and trained to sit in a small primate chair while performing several basic visual tasks to calibrate their eye position (Nummela et al., 2017).

### Eye tracking

Eye position was acquired at 240 Hz using an Arrington Eye Tracker and Viewpoint software (Arrington Research). Eye position was collected from infrared light reflected off a dichroic mirror placed at a 45-degree angle on axis in front of the marmoset. Eye position was calibrated at the start of each behavioral session using two different face detection tasks as described in previous work (Nummela et al., 2017; Coop et al., 2024; Singh et al., 2025). In brief, a set of 3-6 small (1 degree diameter) marmoset faces were placed at random positions on a video display while the marmoset freely viewed them and the experimenter adjusted the center and horizontal/vertical gains of the eye tracking to match eye positions to overlay the objects of interest. After calibration marmosets freely viewed image and video stimuli of the main experimental tasks.

The resulting eye-position data were analyzed using a procedure adapted from a previous study by Coop et al. (2024). Raw horizontal and vertical eye position signals were smoothed offline by convolving with a Gaussian kernel (sigma = 5ms). Saccadic eye movements were detected offline using automatic detection of deviations in 2-D eye velocity space. Horizontal and vertical velocity were computed by taking the difference of smooth eye position and then marking saccades where 2-D velocity greatly exceeded 10 times the median velocity and merging any saccadic events that were within 5 ms in time. Saccade onset and offset were determined by crossing of 2-D median velocity threshold.

Eye traces were collected for the duration of each eye-tracking session, with the marmosets head fixed throughout. Every session began with the calibration of eye traces adapted from previous studies (Nummela et al., 2017; Coop et al., 2024). Marmosets were calibrated using a multi-step routine. They were first presented with circular images of marmoset faces in various preset configurations on a grey background (Fig. 1a). They were allowed to freely view the faces while the locations of their eye traces and the fixation points were plotted. Between trials, vertical/horizontal position and gain of the eye traces were adjusted to be aligned to overlap the position of faces fixated during free viewing. Juice was given at the end of each trial regardless of the monkey’s activity to keep them engaged in the task. Calibration trials continued until the marmoset’s eye traces were aligned as determined by the experimenter. Marmosets then began a new calibration task in which the marmoset faces appeared at randomly sampled locations across the screen (Fig. 1b). In this case marmosets were rewarded if they fixated the faces; if successful, the face was replaced with a fixation point (0.3 degrees diameter), and if they held fixation for 500 ms they were rewarded with juice, at which time the point would disappear encouraging them to sample other points. Eye position and gain were then adjusted as needed using the more stable eye position on fixation points. Again, calibration continued until eye traces were reliably aligned with the fixation targets.

**Figure 1.**
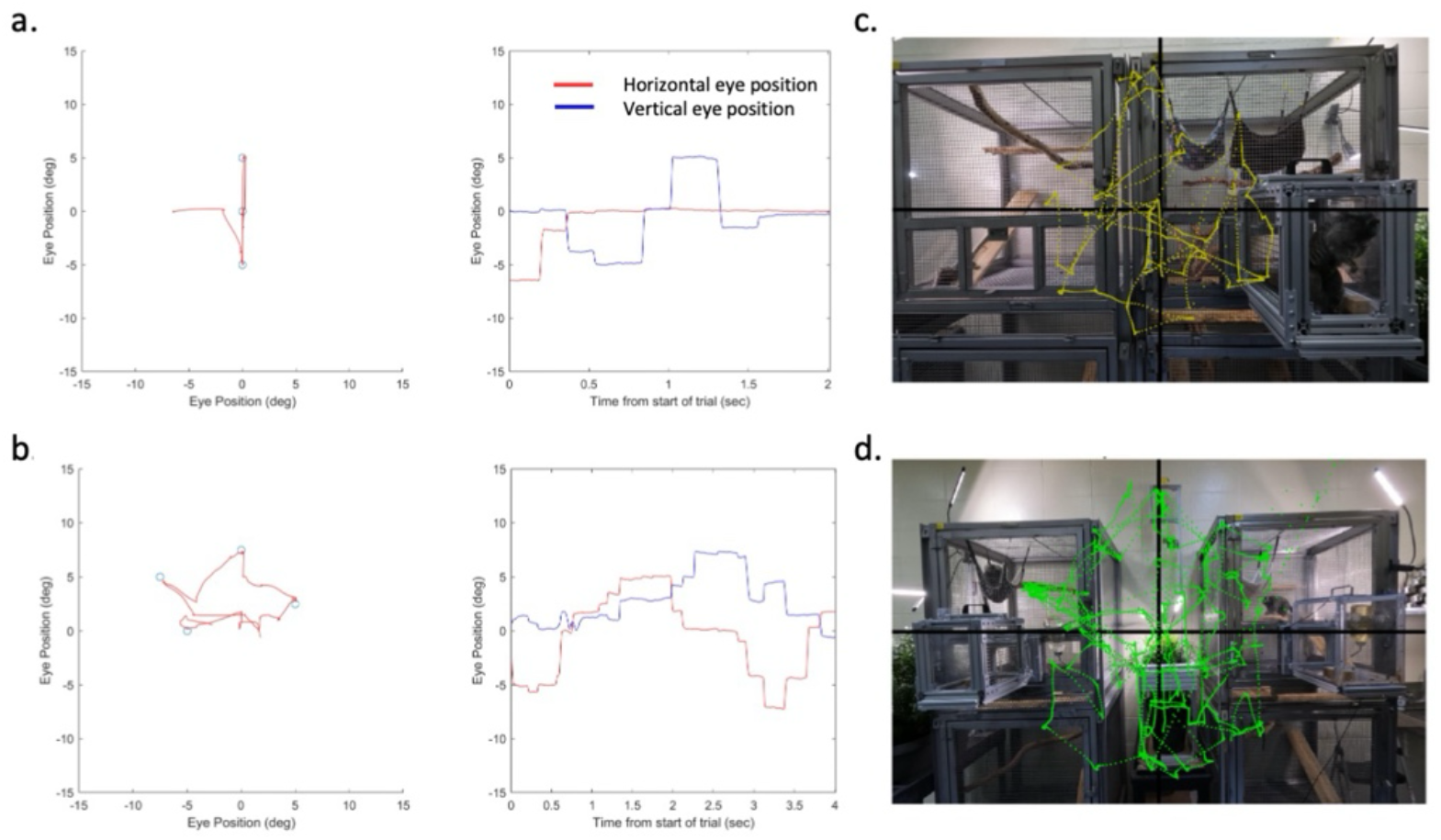
Eye calibration on fixed points and resulting eye position traces and scan paths. (a) Example of the first set of calibration trials, left; blue circles represent the position of marmoset faces and red lines indicate superimposed 2D eye position, right; horizontal (red) and vertical (blue) eye position is shown as function of time from same example trial. Similarly (b) shows example trial from second set of calibration trials in which several faces were positioned randomly and replaced with fixation points upon fixation for reward (left), and corresponding eye position over time (right). (c) Example trial from the task of viewing a static image from the marmoset colony (Monkey A, near condition) with 2D eye traces (yellow) overlaid on image (c) and from a task trial viewing a video with 2D eye traces (green) overlaid on a still frame from video (d). Cage positions in colony video were arranged towards the periphery to encourage viewing animals across the full field of view.

After successful calibration, marmosets began the behavioral task that involved either viewing static natural images or videos sampled from their home marmoset colony. Example 2D eye position traces from the task are shown overlaid on a static image (Fig. 1c) and on a still frame of video (Fig. 1d). To assess the marmosets’ oculomotor range, we used the eye position across trials to compute the normalized density of fixations as a function of radial distance from the center of the screen for both the static images and videos. Histograms of fixation density as a function eccentricity were created from 0-32 dva in bin sizes of 2 dva. Within each 2D bin, we computed the proportion of total fixations in that bin along with confidence intervals using the binofit() function (MATLAB Statistics Toolbox). We also computed the fixation density as a function of 2D x-y space using bins of size 2x2 dva sampled from +/-32 dva to cover the full extent of the screen.

### Stimulus presentation and timing

Stimuli were presented on a gamma-corrected display (BenQ X2411z LED monitor, resolution: 1920 x 1,080, gamma correction 2.2) that had a dynamic luminance range of 0.5-230 cd/m^2^ at 120 Hz. Brightness and contrast were set to 100 and 50, respectively. Additional monitor features were turned off. Gamma corrections were verified using a photometer.

Stimuli were composed of salient videos of marmosets interacting within the home colony of the pair of animals under study. A total of 10 videos were created with a duration of 20 seconds each. Videos were recorded with a GoPro Hero10 camera at 30 frames per second, with a resolution of 1920x1080 pixels, and in video presentation were downsampled to update every 4^th^ frame at 120hz to show them their native speed of 30hz. The content of the videos was intended to mimic what naturally occurs within the home colony, including the presence of caging, colony enrichments and plants, and more so, marmosets interacting. Cages within the colony were moved and configured to create video frames that would contain objects of high interest (moving and interacting conspecifics) spanning across the entirety of the video field of view. This was to promote marmoset gaze in the experiments to reach out to the periphery and to avoid the center bias typical in photography.

We also wanted to ensure salient moving stimuli were present uniformly across a wide range of eccentricities. To confirm our videos accomplished this, we computed motion density maps (Fig. 2). The motion density maps were calculated by summing for each x-y pixel any changes in absolute pixel value across video frames. The average motion density was used to verify that the presented stimuli contained animals moving at appropriate eccentricities from the center, greater than 10 dva. Motion density maps confirmed that our videos have objects of high interest out to around 20 dva (Fig. 2b, left). In a second condition, described below, we presented the stimuli at a closer viewing distance, so the monitor spanned a wider visual field, in which case the motion density was roughly uniform even beyond 25 dva (Fig. 2b, right). This confirmed that the stimuli used in both conditions contained moving objects across a wide range of the screen and would thus be sufficient in promoting eye movements beyond the center of the visual field.

**Figure 2.**
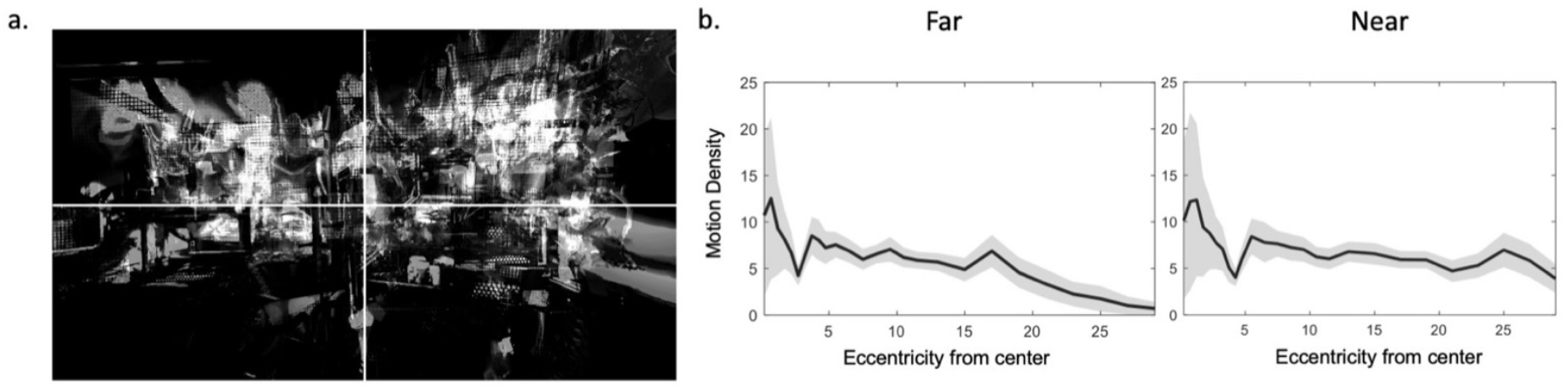
Motion density sampled positions roughly uniformly into the periphery. The stimulus set consisted of a total of 10 videos. To ensure that objects of salience – moving marmosets exhibiting their natural behaviors in the colony – covered a wide range on the screen, the motion density (change in pixel across frames) was averaged across all videos. (a) The superposition of motion density across the 10 videos. (b) Plotting this motion density as a function of eccentricity from center shows that moving marmosets are present on screen out to around 20 dva on screen in the far condition (b, left) and beyond 25 dva on the screen in the near condition (b, right).

Each task trial began with a static image presented for 10 seconds and then followed by a video for 20 seconds (Fig. 3). Static images were randomly drawn still frames from any one of the 10 stimulus videos. After 10 seconds, one of the randomly selected 20-second videos was presented. Marmosets then freely viewed both the static images and the video. In order to ensure that there was enough visual information spanning outside of the previously claimed oculomotor range, both a far and a near condition were tested. In the far condition, marmosets sat at a distance of 57 cm from the screen. This resulted in both the static image and videos spanning ± 25 dva horizontally and ±15 dva vertically from the center of the screen (Fig. 3a). In the near condition, marmosets sat 38 cm away from the screen, resulting in a span of ± 35 dva horizontally and ± 20 dva vertically from the center (Fig. 3b). In both conditions, the behavioral task remained the same: freely viewing static images and the video from the colony. Juice was given every 5 seconds regardless of the monkey’s activity to keep them active during the task. Marmoset A completed a total of 139 trials in the far condition and 50 trials in the near condition, and Marmoset B completed a total of 98 trials in the far condition and 251 trials in the near condition.

**Figure 3.**
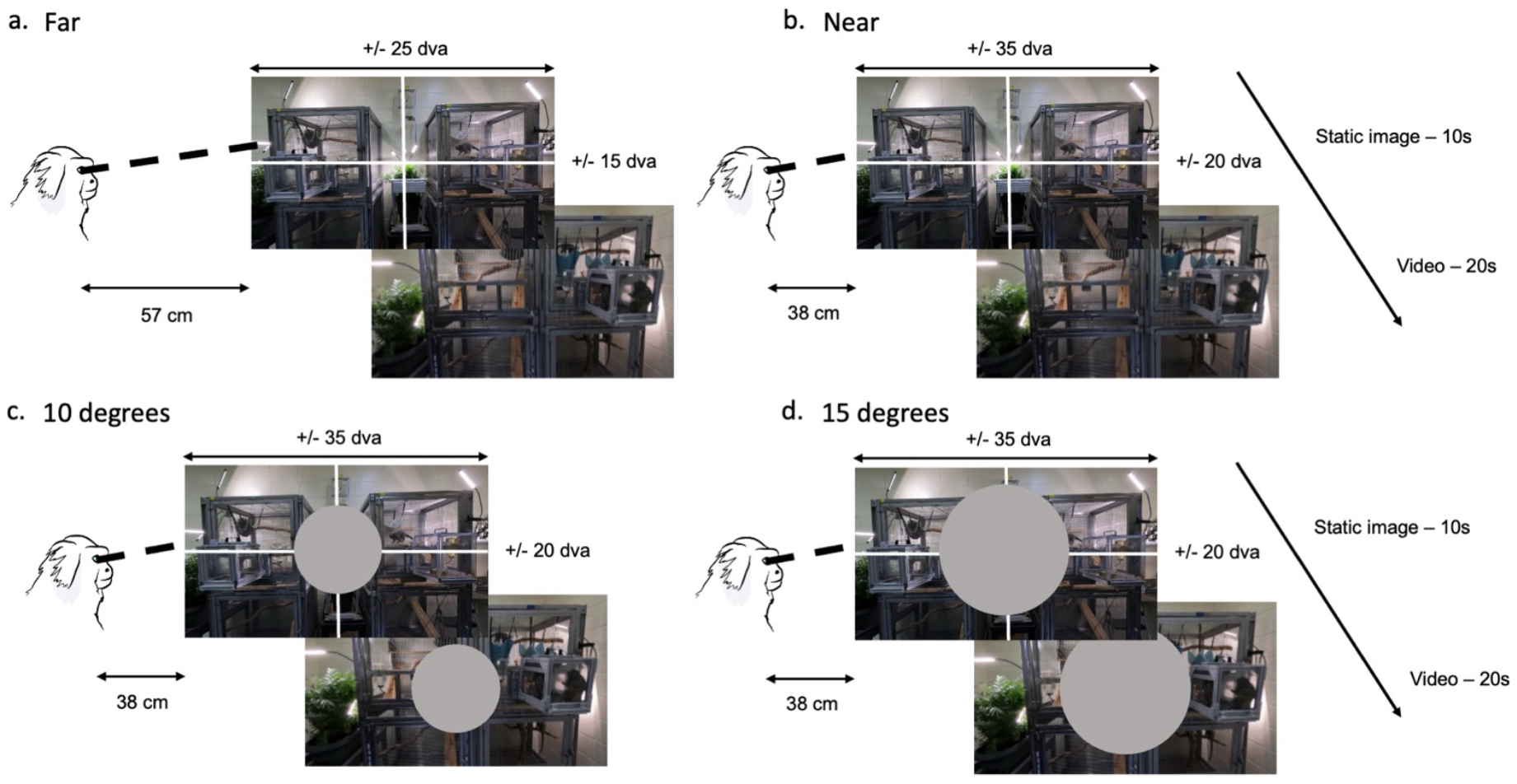
Free-viewing of image and video for far and near viewing, and central occlusion conditions. Each trial consisted of a still image (a randomly selected frame from the video) being presented for 10 seconds followed by a 20-second long video from the colony with freely behaving marmoset monkeys. In the far condition (a) the head fixed marmoset was 57 cm away from the screen resulting in the image and video covering +/-25 dva (degrees visual angle) on the horizontal and +/-15 dva on the vertical. In the near condition (b) the marmoset was 38 cm away from the screen resulting in the image and video covering +/-35 dva on the horizontal and +/-20 dva on the vertical. (c,d) **Central occlusion conditions:** The 10-degree central occlusion condition had a gray circle of radius 10 dva drawn on the center of both the image and video stimuli (c) and the 15-degree central occlusion condition had a circle of radius 15 dva (d). Both central occlusion conditions were presented to subjects in the near viewing conditions (38 cm away).

A central occlusion condition was introduced in one monkey (Marmoset B) to determine if we could encourage marmosets to look more peripherally by removing objects of interest from central vision. In this condition, gray circles of 10 dva (Fig. 3c) and 15 dva (Fig. 3d) were added to the center of the field of view, occluding part of the image or video that remained behind it. All trials were conducted in the near viewing distance (38cm) to maximize the display size. Marmoset B completed 50 trials using the 10 dva occlusion and 89 trials using the 15 dva occlusion.

## Results

In order to determine marmoset oculomotor range, we analyzed normalized fixation density as a function of eccentricity from the center of the screen for both the static images and videos (Fig. 4). To test whether video stimuli resulted in a larger range of eye movements compared to static images, we compared the fixation density distribution for static images with that of the videos. In the far condition, the fixation density distributions for both Marmoset A (Fig. 4a) and Marmoset B (Fig. 4b) did not differ significantly between static images and videos as indicated by the highly overlapping confidence intervals of these distributions. The mean density from averaging the two distributions is shown by the black dashed line overlaid on the distributions to facilitate comparison that they are overlapping. To verify our visual observations, comparison of distributions was performed using a Kolmogorov-Smirnov test. We observed no significant differences in the distribution of fixation density between the static images and videos (across subjects and across the far and near conditions, D = 0.07, p = 0.12, N_*Static*_ = 538, N_*Video*_ = 538). A Wilcoxon rank-sum test on the median fixation eccentricity also showed that this did not differ significantly for static images (M = 7.32 dva) from that of videos (M = 7.35 dva; W-value = 0.07, p = 0.946).

**Figure 4.**
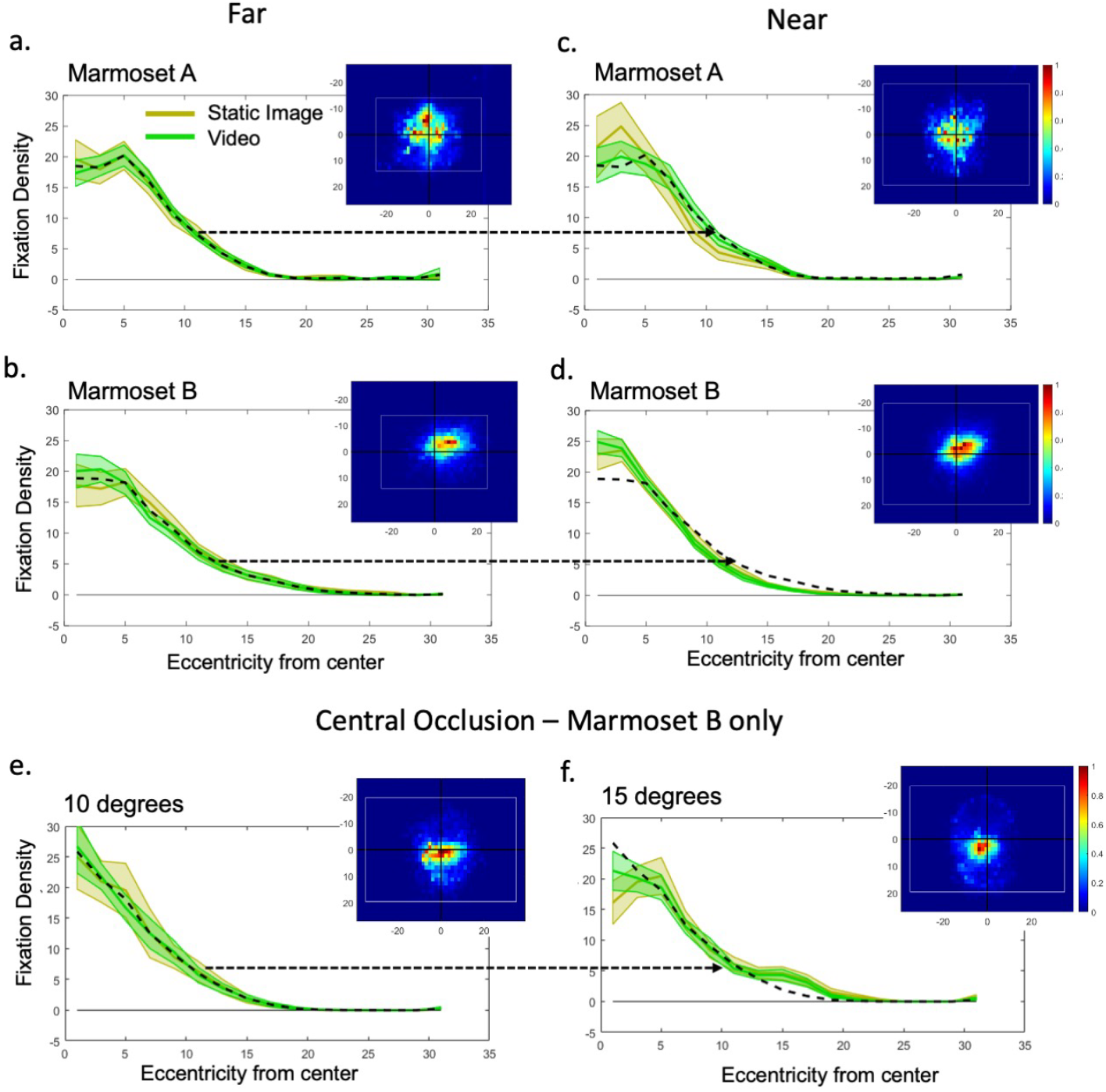
Normalized density of fixations as a function of eccentricity for (a) Marmoset A in the far condition; 139 trials (b) Marmoset B in far condition; 98 trials (yellow for static image and green for video stimuli, shaded regions indicate 2 SEM). Black dotted line in fixation density plots indicates mean fixation density averaged across static image and video for visual comparison. Inset figures show 2D fixation density with white rectangles representing image/video size in dva. Same data for (c) Marmoset A in near condition; 50 trials and (d) Marmoset B in near condition; 251 trials. Black dotted line in fixation density plots indicates mean fixation density for far trials, transposed onto near trials for comparison (indicated by black arrow). For the central occlusion condition (e) shows data from Marmoset B with 10-degree occlusion; 50 trials and (f) also from Marmoset B with 15-degree occlusion; 89 trials. Both were done in the near condition. Black dotted line in fixation density plots indicates mean fixation density for 10-degree occlusion trials, transposed onto 15-degree occlusion trials for comparison (indicated by black arrow).

To encourage marmosets to view more peripherally we included both near and far viewing conditions that had different spans of visual information across the screen. For the near condition moving conspecifics appear even further in the periphery, and thus we might expect if marmoset tracked conspecifics their fixation density would shift peripherally in proportion to the expanded field of view. To confirm this, normalized fixation density distributions with confidence intervals for the static images and videos in the near condition were analyzed. The mean fixation density for both the static images and videos in the far condition as shown by the black dashed line was copied over onto the near condition’s distribution as a visual comparison. Both Marmoset A (Fig. 4c) and Marmoset B (Fig. 4d) show similar distribution shapes between the two conditions, but, contrary to our expectations, with a difference of slightly narrower distributions in the near condition. To quantify this difference, again a two-sample Kolmogorov-Smirnov test was used to compare the distributions. We observed significant differences in fixation density between the far and near conditions (across subjects and across static images and videos, D = 0.28, p < 0.001, N_*Far*_ = 602, N_*Near*_ = 474) and contrary to our expectation, a Wilcoxon rank-sum test showed that the median of the distribution for the near condition (M=6.77 dva) was narrower than for the far condition (M = 7.84 dva; W-value = 8.24, p < 0.001). Thus, placing image and video further into the periphery did not increase the median fixation eccentricity, and even showed a slight decrease. Overall, both Marmoset A and Marmoset B, on average, had a normalized fixation density that remained largely within 15 dva for both the static images and videos, with 70.31% of the fixations occurring within 10 dva and 93.94% occurring within 15 dva. These results are in line with the previous study using natural viewing of static images drawn from the internet (Mitchell et al, 2014).

Although our stimuli provided salient information across the entirety of the screen, a question remained as to whether or not marmosets were physically incapable of looking out further into the periphery or if they could be motivated to do so by only placing stimuli of interest in the periphery. Our central occlusion condition, conducted in the near condition to ensure sufficient objects of interest were in the periphery, addressed this question directly. However, despite these modification to the stimulus, similar results as above were found with the 10 dva central occlusions (Fig. 4e), with 70.35% of the fixations occurring within 10 dva, and 92.16% of all fixations occurring within 15 dva. With the 15-degree occlusion (Fig. 4f), although slight increases in the median fixation density were observed around 7 and 15 dva from the center, fixations still remained largely within 15 dva, with 59.85% of the fixations occurring within 10 dva and 81.73% of all fixations occurring within 15 dva. A two-sample Kolmogorov-Smirnov test on trial-level averages showed no significant difference in fixation densities between static images and videos for both the 10-degree occlusion (D = 0.12, p = 0.869, N = 50) and the 15-degree occlusion (D = 0.12, p = 0.507, N = 89). However, a significant difference was observed between the density of fixations with a 10-degree and 15-degree occlusion (across static images and video, D = 0.29, p < 0.001, N_10-deg_ = 100 and N_15-deg_ = 178). A Wilcoxon rank-sum test on the medians also showed that the median of the distribution with15-degree occlusion (M = 8.37 dva) was significantly higher than with the 10-degree occlusion (M = 7.23 dva; W-value = 3.75 p < 0.001), thus confirming some shift towards more peripheral viewing with the larger central occlusion. Further, both occlusion conditions showed an increase relative to the matching (near) non-occlusion condition (M = 6.77 dva). The 15-degree occlusion also promoted making eye movements more peripherally than the 10-degree occlusion, although the relative increase was not very large (15.7% increase, M = 7.23 dva versus 8.37 dva). Thus, while the occlusions did promote peripheral viewing, there was actually little change in the overall fixation distribution, resulting in marmosets looking mainly at grey regions of screen in these trials.

## Discussion

Humans and non-human primates rely on both eye and head movements to visually scan and gather critical information from the environment. Previous work in smaller primates like the common marmoset and other New World Monkeys has shown limited oculomotor range and larger reliance on head movements to orient attention (J. Burkart & Heschl, 2006; J. M. Burkart & Heschl, 2007; Heiney & Blazquez, 2011; McCrea & Gdowski, 2003; Pandey et al., 2020). Marmosets presented with natural images from the internet in a head-fixed eye tracker exhibited a limited oculomotor range of around 10 dva while scanning the images, leading to open questions about whether this is a physical constraint or a lack of objects of interest to promote eye movements beyond this range. To address this, we employed a head-fixed, free-viewing paradigm using salient naturalistic images and videos as stimuli, which included moving conspecifics, designed to draw the marmosets’ attention to the periphery (up to 25 dva in the far condition and up to 35 dva in the near condition). In addition, we included central gray occluding regions over the images and video to push them to their physical oculomotor limits. Motion density maps confirmed that salient objects (moving conspecifics) spanned the entire range and were not confined to central locations.

Our results show that even under conditions of high peripheral salience, the marmosets’ eye movements remain largely restricted. Across all conditions and animals, around 70% of the fixations were concentrated within 10 dva (in line with previous work, Mitchell et al., 2014) and around 94% of them occurring within 15 dva. There was no statistical difference between fixation density across our two stimulus types, static images and video. Furthermore, when we included a near viewing condition that pushed objects of interest even further into the periphery, we did not observe an increase in fixation density to more peripheral locations – in fact it was even modestly reduced. More so, we included occlusion conditions in which central portion of the stimuli were grayed out, either with a radius of 10 or 15 degrees, and while median eccentricity of fixations did significantly increase in these conditions, the changes were modest. These results together suggest that the limitations may not be due to a lack of salient peripheral stimuli but rather a physiological constraint creating a strong central bias. These findings also underscore that marmosets would more likely use head movement to orient their attention when allowed to do so, which is confirmed in recent head-mounted eye tracking from freely moving marmosets (Singh et al, 2025).

Our findings strongly support the assumptions made in recent marker-free head-tracking studies of freely-moving marmosets (e.g., Meisner et al., 2025; Xing et al., 2024) where the marmoset’s gaze is measured as a 10-degree cone based on the position of the head. Data from head-mounted eye trackers on marmosets also support a similar, or even tighter, range in freely moving marmosets (Singh et al., 2025). Quantifying and verifying this range is essential for ensuring the reliability of the head direction as an estimate of gaze for freely moving marmosets in naturalistic settings, a method that is gaining in popularity for studying interactions between conspecifics in this highly social primate species.

## Acknowledgements

This work was supported by NIH EY030998, NIH T32EY007125 and University of Rochester Research Award. We thank Dina-Jo Graf for technical assistance and Jingwen Li for helpful discussions.

## References

Burkart, J., & Heschl, A. (2006). Geometrical gaze following in common marmosets (Callithrix jacchus). Journal of Comparative Psychology (Washington, D.C.: 1983), 120(2), 120–130. 10.1037/0735-7036.120.2.120

Burkart, J. M., & Finkenwirth, C. (2015). Marmosets as model species in neuroscience and evolutionary anthropology. Neuroscience Research, 93, 8–19. 10.1016/j.neures.2014.09.003

Burkart, J. M., & Heschl, A. (2007). Understanding visual access in common marmosets, Callithrix jacchus: Perspective taking or behaviour reading? Animal Behaviour, 73(3), 457–469. 10.1016/j.anbehav.2006.05.019

Coop, S. H., Yates, J. L., & Mitchell, J. F. (2024). Pre-saccadic Neural Enhancements in Marmoset Area MT. The Journal of Neuroscience: The Official Journal of the Society for Neuroscience, 44(4), e2034222023. 10.1523/JNEUROSCI.2034-22.2023

Guitton, D., & Volle, M. (1987). Gaze control in humans: Eye-head coordination during orienting movements to targets within and beyond the oculomotor range. Journal of Neurophysiology, 58(3), 427–459. 10.1152/jn.1987.58.3.427

Heiney, S. A., & Blazquez, P. M. (2011). Behavioral responses of trained squirrel and rhesus monkeys during oculomotor tasks. Experimental Brain Research, 212(3), 409–416. 10.1007/s00221-011-2746-4

Kano, F., & Tomonaga, M. (2009). How chimpanzees look at pictures: A comparative eye-tracking study. Proceedings. Biological Sciences, 276(1664), 1949–1955. 10.1098/rspb.2008.1811

McCrea, R. A., & Gdowski, G. T. (2003). Firing behaviour of squirrel monkey eye movement-related vestibular nucleus neurons during gaze saccades. The Journal of Physiology, 546(Pt 1), 207– 224. 10.1113/jphysiol.2002.027797

Meisner, O. C., Shi, W., Nair, A., Nandy, G., Jadi, M. P., Nandy, A. S., & Chang, S. W. C. (2025). Diverse and flexible strategies enable successful cooperation in marmoset dyads. Current Biology: CB, 35(18), 4509-4521.e5. 10.1016/j.cub.2025.08.005

Miller, C. T., Freiwald, W. A., Leopold, D. A., Mitchell, J. F., Silva, A. C., & Wang, X. (2016). Marmosets: A Neuroscientific Model of Human Social Behavior. Neuron, 90(2), 219–233. 10.1016/j.neuron.2016.03.018

Mitchell, J. F., Reynolds, J. H., & Miller, C. T. (2014). Active vision in marmosets: A model system for visual neuroscience. The Journal of Neuroscience: The Official Journal of the Society for Neuroscience, 34(4), 1183–1194. 10.1523/JNEUROSCI.3899-13.2014

Nummela, S. U., Coop, S. H., Cloherty, S. L., Boisvert, C. J., Leblanc, M., & Mitchell, J. F. (2017). Psychophysical measurement of marmoset acuity and myopia. Developmental Neurobiology, 77(3), 300–313. 10.1002/dneu.22467

Pandey, S., Simhadri, S., & Zhou, Y. (2020). Rapid Head Movements in Common Marmoset Monkeys. iScience, 23(2), 100837. 10.1016/j.isci.2020.100837

Singh, V. P., Li, J., Dawson, K., Mitchell, J. F., & Miller, C. T. (2025). Active vision in freely moving marmosets using head-mounted eye tracking. Proceedings of the National Academy of Sciences, 122(6), e2412954122. 10.1073/pnas.2412954122

Tehovnik, E. J., Froudarakis, E., Scala, F., Smirnakis, S. M., Patel, S. S., & Tolias, A. S. (2021). Visuomotor control in mice and primates. Neuroscience & Biobehavioral Reviews, 130, 185– 200. 10.1016/j.neubiorev.2021.08.009

Xing, F., Sheffield, A. G., Jadi, M. P., Chang, S. W. C., & Nandy, A. S. (2024). 4Dynamic modulation of social gaze by sex and familiarity in marmoset dyads (p. 2024.02.16.580693). bioRxiv. 10.1101/2024.02.16.580693

